# Spatiotemporal dynamics of self-generated imagery reveal a reverse cortical hierarchy from cue-induced imagery

**DOI:** 10.1101/2023.01.25.525474

**Authors:** Yiheng Hu, Qing Yu

## Abstract

Visual imagery, the ability to generate visual experience in the absence of direct external stimulation, allows for the construction of rich internal experience in our mental world. Most imagery studies to date have focused on cue-induced imagery, namely the to-be-imagined contents were triggered by external cues. It has remained unclear how internal experience derives volitionally in the absence of any external cues, and whether this kind of self-generated imagery relies on an analogous cortical network as cue-induced imagery. Here, leveraging a novel self-generated imagery paradigm, we systematically examined the spatiotemporal dynamics of self-generated imagery, by having participants volitionally imagining one of the orientations from a learned pool; and of cue-induced imagery, by having participants imagining line orientations based on associative cues acquired previously. Using electroencephalography (EEG) and functional magnetic resonance imaging (fMRI), in combination with multivariate encoding and decoding approaches, our results revealed largely overlapping neural signatures of cue-induced and self-generated imagery in both EEG and fMRI; yet, these neural signatures displayed substantially differential sensitivities to the two types of imagery: self-generated imagery was supported by an enhanced involvement of anterior cortex in generating and maintaining imagined contents, as evidenced by enhanced neural representations of orientations in sustained potentials in central channels in EEG, and in posterior frontal cortex in fMRI. By contrast, cue-induced imagery was supported by enhanced neural representations of orientations in alpha-band activity in posterior channels in EEG, and in early visual cortex in fMRI. These results jointly support a reverse cortical hierarchy in generating and maintaining imagery contents in self-generated versus externally-cued imagery.

## Introduction

Visual imagery is the ability to generate visual experience from the internal world, in the absence of direct external stimulation ^1^. It remains a fundamental capability of human cognition, and is central to the understanding of how our mental world is constructed. Unlike visual perception which is primarily driven by physical external stimulation and can be measured via standardized paradigms, visual imagery by definition involves cognitive processes that are ambiguous and difficult to measure in nature. Consequently, various behavioral paradigms have been used to study visual imagery; these paradigms might fundamentally differ in the exact cognitive processes involved, yet most of them share a cue-induced nature in common, namely the contents of imagery are induced by externally presented cues. Overall, two types of imagery tasks are most frequently used: the first type of imagery task employs semantic ^2-4^ or associative cues ^5^ to trigger retrieval of imagery contents from long-term memory. In these tasks, the to-be-imagined contents are not directly accessible on the screen, but can only be inferred from long-term memory. The other type of imagery task utilizes retrocues ^6,7^ or mental rotation cues ^8,9^ to access specific memorized contents maintained or manipulated in working memory. These cue-induced imagery tasks, albeit significantly differed in their way to cue imagery, have led to several consistent observations in visual imagery: first, imagery and perception share common neural codes in early visual cortex for simple visual features ^8^, in object-selective high-level visual cortex for complex visual objects ^2,10^. and in alpha-band activity in electroencephalography (EEG) ^4^, suggesting the depictive nature of visual imagery; second, neural processing during imagery follows a reverse cortical hierarchy from that during perception, which is supported by larger spatial overlap of univariate BOLD activations between imagery and perception in higher-order frontoparietal than in occipitotemporal cortex ^3^; an increased top-down signal flow in imagery compared to perception, from frontal ^11,12^ or parietal ^13^ to occipital cortex; and a reversal of object representations from high-level to low-level visual cortex ^2,14,15^. These findings together indicated that visual imagery involves a distributed cortical network from low-level visual cortex to higher-level visual and frontoparietal cortex ^3,16^, and provided empirical support for the reverse visual hierarchy model, which proposes that, as opposed to perception which triggers a feedforward sweep of neural activations along the posterior-to-anterior cortical hierarchy, imagery is initiated by top-down signals generated in higher-level cortex that trigger a cascade of neural processing in the downstream cortical areas eventually ^1,17^.

However, imagery experience by definition can be generated in the absence of any external stimulation, including external cues. In this context, imagery is entirely perception-or cue-independent, and the contents of imagery are self-generated from the internal world. This self-generated imagery can be regarded as a part of self-generated cognitive processes, during which an internal experience arises from intrinsic changes within an individual, rather than extrinsic changes cued from the external environment ^18^. As such, self-generated imagery is much less prone to external influences, and may better reflect “pure” internally-generated mental processes. Although there have been studies on self-generated cognitive processes related to imagery, such as recalling memories, envisioning the future, and mind wandering ^18,19^, those studies engaged complex cognitive processes wherein imagery was only a part of the processes. The neural mechanism of pure self-generated imagery has remained elusive. Specifically, it remains unclear whether self-generated imagery works fundamentally differently from cue-induced imagery, and whether the neural principles with classic cue-induced imagery paradigms would hold with self-generated imagery.

Given that self-generated and cue-induced imagery differ primarily in the origin of imagery contents, it is plausible that when participants orient internally and determine their imagery contents volitionally, the reverse cortical hierarchy might be involved differently from that during cue-induced imagery. A previous study has observed increased decoding performance of imagined objects, as opposed to degraded decoding performance of perceived objects, from low-level to high-level visual cortex ^2^. The rationale is that if one brain region serves as the neural locus that initiates imagery contents signals at the top of the reverse hierarchy, this region should demonstrate better decoding performance of imagined than perceived contents, compared to other downstream brain regions. With this logic, we would expect to see differential representational signals between self-generated and cue-induced imagery along a reverse hierarchy of imagery, due to the internal origin of self-generated imagery.

Here we set out to address these questions by comparing the neural processes underlying self-generated imagery with those of cue-induced imagery. To study self-generated imagery in well-controlled settings and to reduce ambiguity in imagery contents, participants’ imagery contents were constrained to a pool of seven fixed line orientations throughout the experiment. In self-generated imagery, participants determined their imagery content freely without any associative sensory input, one at a time from the seven orientations on each trial; In cue-induced imagery, participants imagined one orientation based on an externally-presented associative cue, and the associations between orientations and cues were learned prior to the task. We investigated the spatiotemporal dynamics of these two types of imagery in a series of two experiments, using EEG (in Experiment 1) and fMRI (in Experiment 2), respectively. In both experiments, neural representations of imagery contents were characterized using inverted encoding models (IEMs), which have been shown to be a powerful tool in unveiling population-level, feature-selective representations across visual, parietal, and frontal cortex in the visual working memory literature ^20-22^.

To preview, across two experiments, we demonstrated self-generated and cue-induced imagery shared common neural representations within multiple neural signatures, while preserving substantial differences in terms of the strength of representations at different levels of cortical hierarchy: in EEG, enhanced orientation representations were observed in self-generated imagery compared to cue-induced imagery in sustained potentials in central channels, and the opposite was true in alpha-band oscillatory activity in posterior channels. In fMRI, enhanced orientation representations were observed in self-generated imagery in right superior precentral sulcus (sPCS) of frontal cortex, and the reverse was true in early visual cortex (EVC). In other words, the relative representational strength of self-generated and cue-induced imagery also followed a frontal-to-occipital reverse hierarchy. Together, these results provided the first empirical evidence, to our knowledge, that frontal cortex plays a critical role in the generation and maintenance of self-generated imagery contents, supporting and extending the reverse hierarchy theory of imagery.

## Results

### EEG Behavior results

In Experiment 1, participants performed an imagery task along with EEG recording (Figure 1A), during which their imagery contents were either cued by one of seven pairs of learned associations between kaleidoscope images and line orientations (Cue-induced Imagery), or self-generated from the same set of seven orientations (Self-generated Imagery). During the learning session, participants successfully acquired the associations between kaleidoscope images and line orientations with their mean absolute recall errors being below 10°. During the EEG session, participants performed the cue-induced imagery task with a mean absolute recall error of 9.47° (SD = 14.28°).

**Figure 1.**
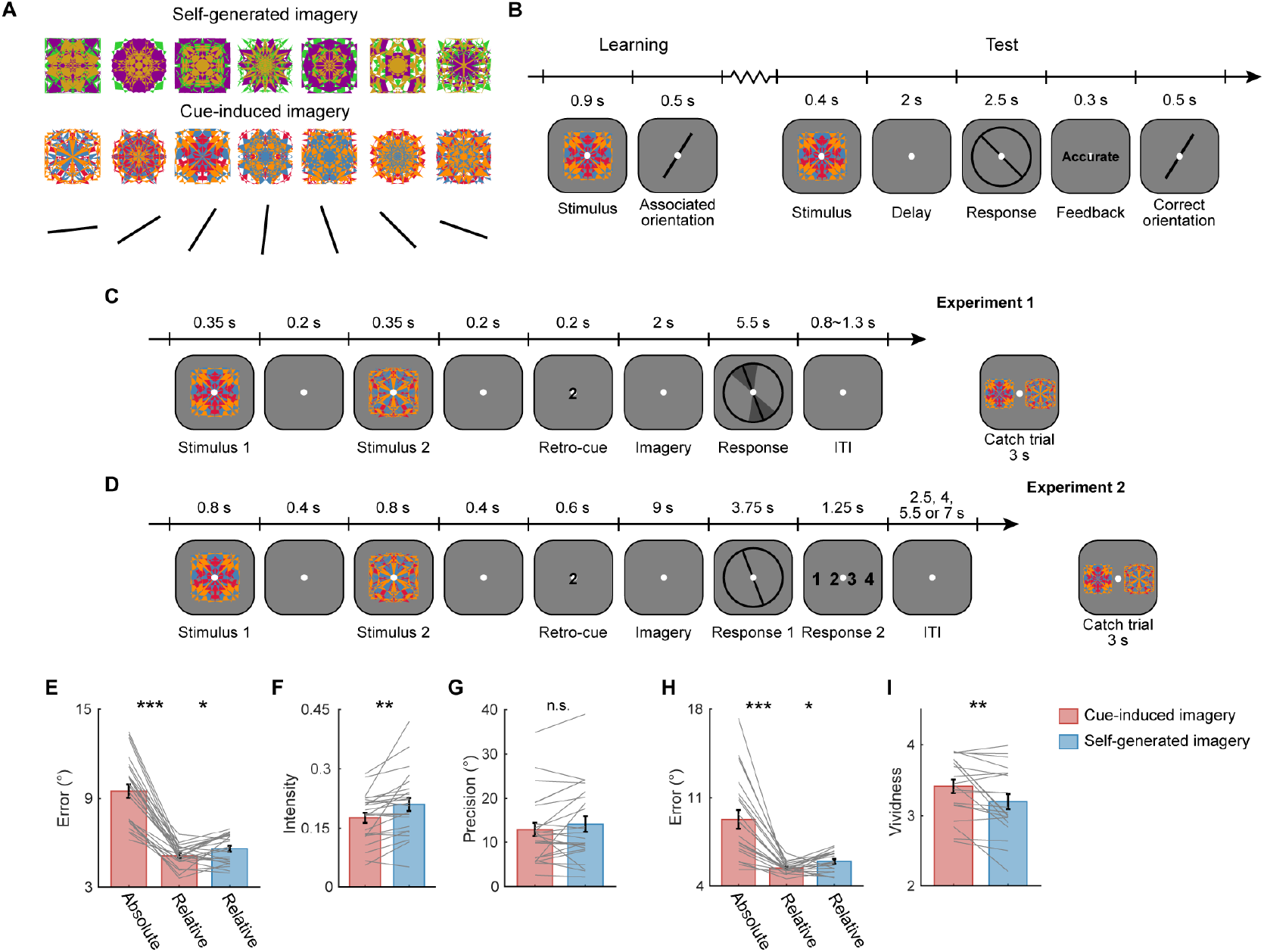
Experimental paradigms and behavioral results. A. Kaleidoscope images and line orientations used in the current study. Two sets of kaleidoscope images were used, each consisted of seven distinct images. The specific set of kaleidoscope images used for each condition (cue-induced or self-generated imagery) was counterbalanced across participants. The specific association between each kaleidoscope image and each orientation was also randomized across participants. B. Trial structure of learning and test tasks. On learning trials, participants passively viewed one kaleidoscope image, followed by its associated line orientation. On test trials, participants viewed one kaleidoscope image, and were required to report its associated orientation. Feedback was provided at the end of each trial. C-D. Trial structure of the main task. A similar trial structure was used in Experiments 1 (C) and 2 (D), and only the timing of events and the type of responses differed. Each trial began with the presentation of two consecutive kaleidoscope images followed by a retrocue. In cue-induced imagery, participants actively imagined the line orientation associated with the cued kaleidoscope image during delay; in self-generated imagery, participants freely chose one from the seven learned orientations and imagined the self-generated orientation during delay. In Experiment 1, participants reported the imagined orientation, the precision, and the intensity of their imagery; In Experiment 2, participants reported the imagined orientation and 1-4 points of vividness rating. Catch trials were interleaved to maintain participants’ attention on the kaleidoscope images, and participants needed to choose the cued kaleidoscope from two probe images after retrocue. E-I. Behavioral performance in Experiments 1 and 2. E. Results of mean recall error in each condition (absolute recall error in cue-induced imagery, and relative recall error in both conditions) of Experiment 1. Colored bars indicate group mean (error bars denote ±1 SEM), gray lines indicated results from individual participants. Asterisks on top denote significance of pairwise comparisons between conditions, n.s., not significant, *: *p* < 0.05, **: *p* < 0.01, ***: *p* < 0.001. F. Same as E, but with results of intensity of imagery experience in Experiment 1. G. Same as F, but with results of precision of imagery experience in Experiment 1. H. Same as E, but with results from Experiment 2. I. Same as E, but with results of vividness rating of Experiment 2.

Because there were no correct answers on self-generated imagery trials, to compare the behavioral performance between conditions, we took the least circular distance of responses to the seven specific orientations as relative recall errors in both conditions. We first showed that relative and absolute errors correlated with each other in cue-induced imagery (*r* = 0.54, *p* = 0.006; Figure S1A), suggesting relative error may be treated as an approximation of absolute error when the latter was not available in self-generated imagery. Meanwhile, relative error was significantly smaller than absolute error in cue-induced imagery, *t*(23) = 11.25, *p* < 0.001. When comparing relative error between conditions, we found that the mean relative error in cue-induced imagery (5.11° ± 3.58°) was slightly but significantly smaller than that in self-generated imagery (5.58° ± 3.72°), *t*(23) = 2.49, *p* = 0.020 (Figure 1E). Furthermore, because participants were required to randomly select one from seven learned orientations in self-generated imagery, we examined whether participants’ responses were biased towards specific orientation bins. We binned all responses into seven bins, each centered at one of the seven orientations (Figure S2A, S2B). We observed a slight bias in participants’ response distribution in both conditions. To avoid potential influence of these biases on subsequent neural analyses, we balanced the number of trials within each response bin for all neural analyses (see Methods for details). Lastly, we confirmed that participants did not respond by simply entering the initial orientation of the response wheel (Figure S2C, S2D). Together, these results suggested that participants faithfully followed task instructions and randomly selected one from seven orientations in self-generated imagery.

Besides recall errors, participants were also measured on the vividness of their imagery, by reporting both the precision (as characterized by the angle of the response wedge) and the intensity (as characterized by the darkness of the response wedge) of their imagery experience. Overall, participants’ subjective experience was more vivid in cue-induced imagery than in self-generated imagery: participants reported a more intense imagery experience in cue-induced imagery (0.18 ± 0.13) compared to in self-generated imagery (0.21 ± 0.14) condition, *t*(23) = 3.37, *p* = 0.003 (Figure 1F). On the contrary, difference in precision between conditions was numerically but not statistically different (12.96° ± 11.47° in cue-induced imagery; 14.24° ± 12.48° in self-generated imagery; *t*(23) = 1.43, *p* = 0.17; Figure 1G). These results indicated that self-generated imagery produced an attenuation of subjective experience in terms of subjective intensity, but not in subjective precision. Moreover, in self-generated imagery, precision significantly correlated with intensity (*r* = 0.58, *p* = 0.003, Figure S1C) and relative errors (*r* = 0.40, *p* = 0.050, Figure S1D) across participants, while the correlation between intensity and relative errors was not significant (*r* = 0.07, *p* = 0.747, Figure S1H). Follow-up stepwise regression analysis confirmed that intensity and relative error explained distinct variance in precision (*p*s = 0.002 and 0.032, respectively). In comparison, no correlation was observed between any two of the behavioral measures in cue-induced imagery (*r*s< 0.40, *p*s > 0.05).

### Sustained potentials and alpha-band oscillatory activity showed differential sensitivity to self-generated and cue-induced imagery

Having established that participants could faithfully perform the self-generated imagery task following task instructions, we next seek to investigate neural signals that could potentially distinguish self-generated from cue-induced imagery. For this purpose, we chose to focus on stimulus-specific neural representations of imagery contents (i.e., orientations in the current study). Specifically, we used participants’ responses on each trial to reconstruct population-level, orientation-selective representations from EEG signals using multivariate inverted encoding models (IEMs). This approach has been successfully applied to investigate orientation representations in various cognitive functions, for both maintenance in working memory ^20,21^ and retrieval from long-term memory ^23^. Previous studies have successfully decoded imagery contents from alpha-band ^4^ as well as voltage signals ^6^ in EEG. On the other hand, recent studies on working memory ^24,25^ indicated that alpha power and sustained potentials in EEG might reflect signals from distinct cognitive processes. Given the close link between imagery and working memory ^8^, here we performed IEM analyses on both voltage and alpha-band (8 – 12 Hz) oscillatory signals (Figure 2A), to investigate whether voltage and oscillatory signals played differential roles in self-generated and cue-induced imagery.

**Figure 2.**
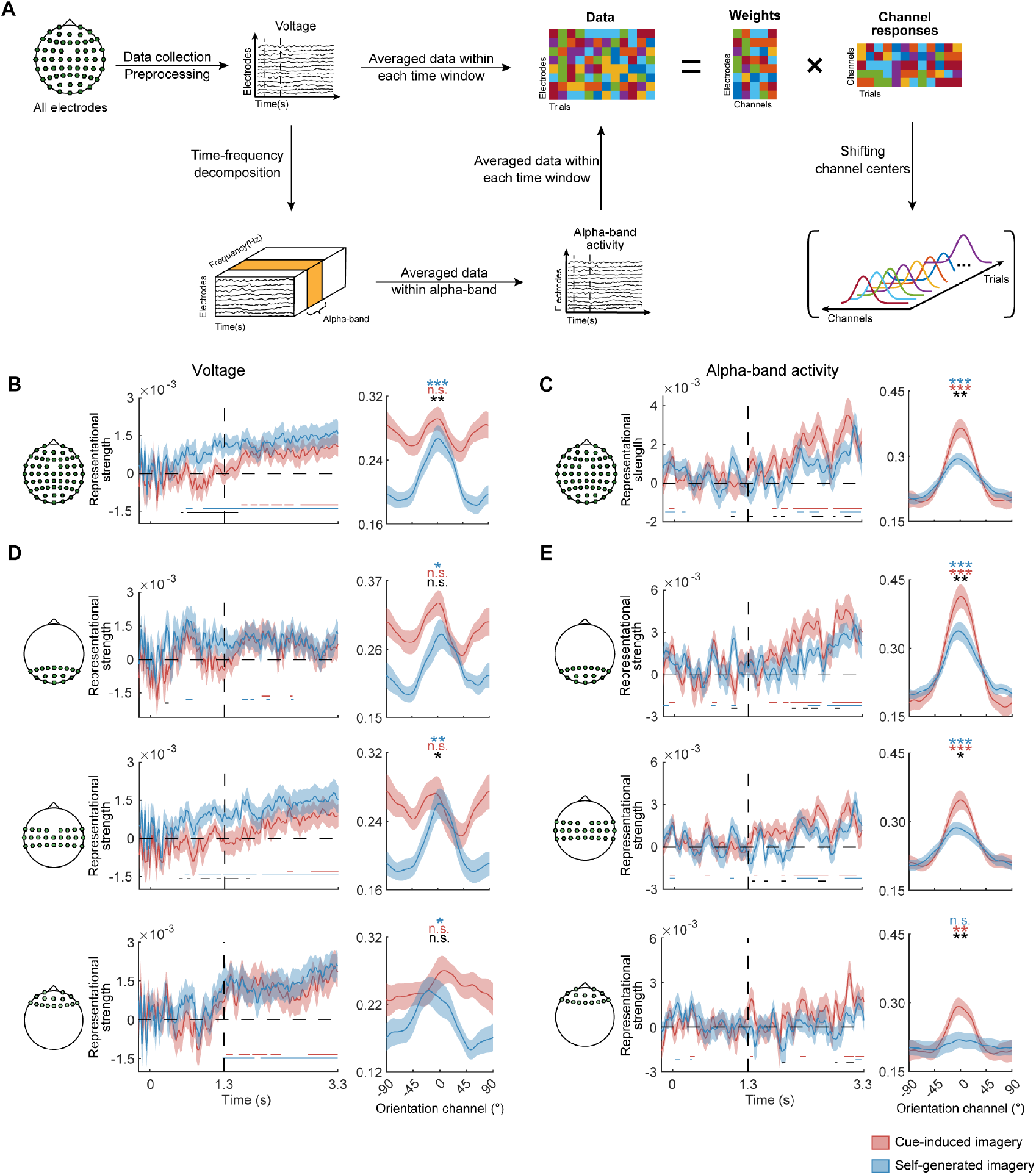
EEG analysis pipeline and results. A. Pipeline of EEG analyses. Raw EEG data were collected for all electrodes. After preprocessing, preprocessed voltage data were fed into IEM analyses. Alternatively, preprocessed voltage data underwent time frequency decomposition, and the obtained power data of different frequency bands were fed into IEM analyses. B. IEM results from voltage data in all electrodes. The left panel shows time course of the strength of orientation reconstructions in cue-induced (red) and self-generated imagery (blue), from -0.2 s prior to stimulus onset until end of delay. Y axis denotes orientation representational strength, quantified using the slope of orientation reconstructions. Colored lines at the bottom denote significant time points of the corresponding condition, corrected for multiple comparisons using a cluster-based permutation method (*p* < 0.01). The vertical dashed line denotes onset of delay (at 1.3 s). The horizontal dashed line denotes baseline of reconstructions. Shaded areas denote error bars (±1 SEM). The right panel shows orientation reconstructions averaged over the selected time period of significance (0.6 – 1.7 s), in cue-induced (red) and self-generated imagery (blue). X axis represents distance from response orientations, with 0 representing the response orientation of each trial. Y axis represents reconstructed orientation channel responses in arbitrary units. Colored asterisks denote significance of the corresponding condition, and black asterisk denotes significance of difference between conditions. n.s., not significant, *: *p* < 0.05, **: *p* < 0.01, ***: *p* < 0.001. C. same as B, but with results from alpha-band power data in all electrodes, and orientation reconstructions were computed over a time window of 2-3.3 s. D. IEM results from voltage data in posterior (top), central (middle), and frontal (bottom) electrodes, using the same analyses and illustrations as in B. E. IEM results from alpha-band power data in posterior (top), central (middle), and frontal (bottom) electrodes, using the same analyses and illustrations as in C.

Our results demonstrated that imagined orientations were represented in both voltage and oscillatory signals during memory delay. Interestingly, the temporal evolution of imagery representations significantly differed in these two types of signals: in voltage signals, significant representations of self-generated imagined orientations ramped up around retrocue period (0.6 s after trial onset) and sustained till the end of delay; whereas significant representations of cue-induced imagined orientations emerged later in time and was much less stable (Figure 2B; all results reported here and in subsequent analyses were corrected for multiple comparisons using a cluster-based permutation method). We quantified this difference by comparing the representational strength of self-generated imagery and that of cue-induced imagery during a temporal epoch around retrocue (0.6 – 1.7 s after trial onset): the representational strength of orientations in self-generated imagery was significant, *t*(23) = 3.97, *p* = 0.0003, and was significantly higher than that in cued-induced imagery, *t*(23) = 3.15, *p* = 0.002. Meanwhile, the representational strength of orientations in cue-induced imagery did not reach significance, *t*(23) = 0.14, *p* = 0.447. By contrast, in alpha-band activity, significant and stable representations of both self-generated and cue-induced imagined orientations ramped up around midway into the delay (2 s after trial onset; Figure 2C). Moreover, when comparing the representational strength of imagery between conditions, a reversed pattern was observed during late delay (2 - 3.3s): the representational strength of orientations in self-generated imagery was significantly weaker than that in cue-induced imagery, *t*(23) = 3.18, *p* = 0.002. To validate the opposite results in voltage and oscillatory signals, we sorted participants’ responses into seven bins and performed multi-class classification on binned orientations using support vector machines (SVMs). Representational differences between conditions in both alpha-band and voltage signals remained during late delay (2 - 3.3s) with the decoding approach (Figure S3). This result confirmed that the observed pattern was robust across different analytical approaches used to reveal orientation representations.

To examine the spatial configuration of electrodes that might have contributed to the representational differences between conditions, we restricted the IEM analyses to frontal, central and posterior EEG electrodes, respectively. We found that, for voltage signals, only central electrodes showed earlier emergence of self-generated representations, and stronger orientation representations in self-generated than in cue-induced imagery (*t*(23) = 2.31, *p* = 0.015; Figure 2D), suggesting that self-generated representations might primarily derive from central electrodes activity. In frontal and posterior electrodes, only self-generated imagery demonstrated weak orientation representations, and no difference remained in terms of either temporal dynamics or representational strength between conditions (frontal: *t*(23) = 1.97, *p* = 0.030 in self-generated imagery, *t*(23) = 0.80, *p* = 0.217 in difference; posterior: *t*(23) = 2.12, *p* = 0.022 in self-generated imagery, *t*(23) = 1.13, *p* = 0.134 in difference). In alpha-band activity, the representational strength of orientations in cue-induced imagery was higher than that in self-generated imagery in posterior electrodes (*t*(23) = 2.96, *p* = 0.004; Figure 2E). Similar but weaker patterns were observed in central (*t*(23) = 2.21, *p* = 0.018) and frontal electrodes (*t*(23) = 3.44, *p* = 0.001). In addition, to examine the specificity of the effect to alpha-band activity, we repeated the analyses across frequencies ranging from 3 to 45 Hz in posterior electrodes. We confirmed that among all frequencies, alpha-band demonstrated the strongest orientation representations, as well as differences between conditions. In addition, similar results were also observed in part of beta- and theta-band, but the effects were overall weaker and less stable (Figure S4).

In Experiment 1, we demonstrated that while self-generated and cue-induced imagery shared representations in both voltage and alpha-band oscillatory signals, the strength of orientation representations carried in these two types of signals significantly differed between conditions in at least two aspects: first, orientation representations in self-generated imagery was stronger than those in cue-induced imagery in voltage signals, and the reverse was true in alpha-band signals. Second, difference in voltage signals was mainly contributed by central electrodes, while difference in oscillatory signals was mainly contributed by posterior electrodes. Because the trial structure of these two conditions were identical, and the only difference was that imagery contents were determined by different sources (self-generated versus externally-cued), we speculated that the differences in signal types and spatial configurations might have reflected differences in internally-generated versus externally-driven imagery: voltage signals from more anterior electrodes might have reflected contents derived from self-generated imagery, and alpha oscillations from more posterior electrodes might have carried information related to externally-driven processing. This anterior versus posterior contrast in the spatial layout of electrodes might reflect a reverse hierarchy in processing information from self-generated versus externally-cued imagery. However, due to the limited spatial resolution of EEG signals, we refined our approach in a second experiment during which we leveraged fMRI to investigate possible neural loci of the reverse hierarchy.

### fMRI Behavior results

In order to identify brain regions that might underlie the representational differences between cue-induced and self-generated imagery in voltage and alpha-band signals, we had participants performed the imagery task inside an MRI scanner in Experiment 2. The procedure of Experiment 2 was similar to that of Experiment 1, except that the timing of events and type of responses were adjusted to better suit for fMRI. Specifically, the delay period was prolonged to compensate for the sluggishness of BOLD signals, and vividness rating of 1-4 points was used in order to shorten the response time inside the scanner (Figure 1D).

The behavioral results in Experiment 2 largely replicated those in Experiment 1: the mean absolute error in cue-induced imagery was 9.30° (SD = 14.15°); the mean relative errors were 5.41° (SD = 3.64°) in cue-induced imagery and 5.94° (SD = 3.71°) in self-generated imagery. Relative errors were significantly smaller in cue-induced compared to self-generated imagery, *t*(19) = 2.78, *p* = 0.012 (Figure 1H). In terms of vividness rating, participants reported a more vivid experience in cue-induced imagery (3.42 ± 0.63) than in self-generated imagery (3.20 ± 0.71), *t*(19) = 2.98, *p* = 0.008 (Figure 1I). Moreover, vividness did not correlate with relative errors in either condition, *r*s < 0.24, *p*s > 0.32. In combination with results from Experiment 1, these results together suggested that precision and intensity likely reflected two different dimensions of vividness, with intensity producing qualitatively similar measures as vividness ratings.

### Whole-brain identification of representations of imagined orientations

To localize brain regions showing differences in representations of imagined orientations between self-generated and cue-induced imagery, we conducted a whole-brain searchlight analysis in combination with IEM. Considering a typical hemodynamic lag of 4-6 s, the searchlight was performed on data of the memory delay period (9 -12 s). Significance of searchlight results were evaluated using one-tailed t-test (*p* < 0.05) and multiple comparison correction (FWE-corrected *p* < 0.01) to obtain the statistical parametric maps.

The whole-brain searchlight revealed largely overlapping brain regions for both cue-induced and self-generated imagery: significant clusters with robust neural representations of imagined orientations were found in a distributed network of cortical regions, including primary visual cortex (V1), extrastriate cortex, intraparietal sulcus (IPS), middle and superior temporal sulcus (STS), left superior precentral sulcus (sPCS), left superior medial gyrus and left middle and inferior frontal gyrus. Besides these common brain regions with shared neural representations, additional clusters were identified separately for the two conditions: in cue-induced imagery, imagined orientations were represented in right inferior frontal sulcus (Figure 3A); in self-generated imagery, imagined orientations were represented in right sPCS and right rostral lateral prefrontal cortex (rlPFC; Figure 3B).

**Figure 3.**
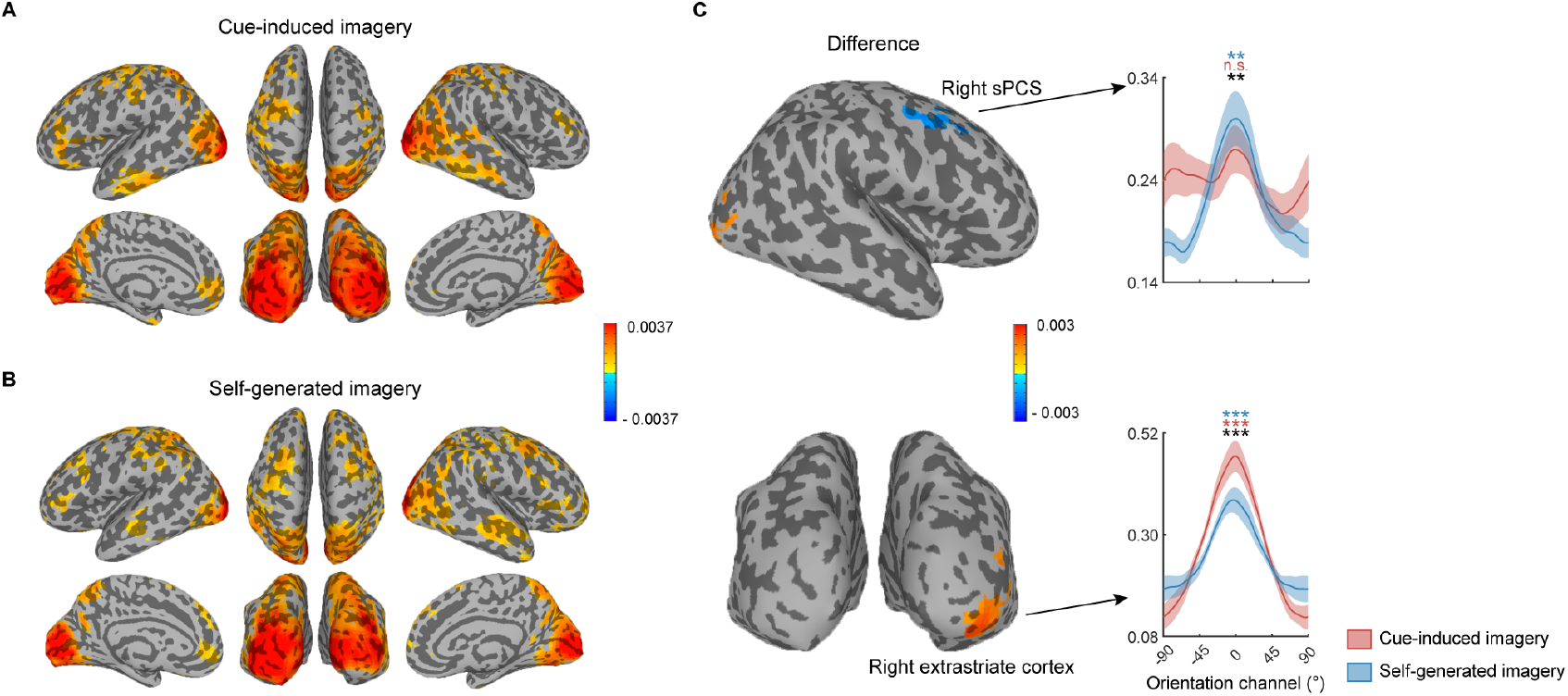
Whole-brain neural representations of imagery contents in late delay. A. Searchlight parametric map of the strength of orientation representations in late delay (9-12 s; 6 s after retrocue) in cue-induced imagery. Colors on the cortical surface denote brain regions with significant orientation representations, corrected using a cluster-based permutation method (*p* < 0.01). For demonstration purposes, clusters were thresholded at 50 voxels. B. Same as A, but with results from self-generated imagery. C. Difference map of orientation representations in A and B, with positive values denoting stronger orientation representations in cue-induced imagery, and negative values denoting stronger orientation representations in self-generated imagery. Orientation reconstructions obtained from the two significant clusters were shown, with right sPCS (top panel) demonstrating stronger orientation representations in self-generated imagery, and right extrastriate cortex demonstrating stronger orientation representations in cue-induced imagery. X axis represents distance from response orientations, with 0 representing the response orientation of each trial. Y axis represents reconstructed orientation channel responses in arbitrary units. Colored asterisks denote significance of cue-induced (red) and self-generated (blue) imagery, and black asterisk denotes significance of difference between conditions. n.s., not significant, *: *p* < 0.05, **: *p* < 0.01, ***: *p* < 0.001.

Previous studies have revealed shared representations of perception and imagery in EVC. In the current study, participants were not exposed to physical line orientations in either condition throughout the imagery task. To investigate the nature of the imagery representations and to verify that participants did engage visual imagery in Experiment 2, we had participants performed a perception task of orientations inside the scanner following the main imagery task. We then trained an IEM with perception data, and tested the perception model on imagery data in a second “perception” searchlight analysis. The perception searchlight revealed similar clusters in visual cortex (Figure S5), confirming a perception-like neural representation of orientations in our imagery task. Additionally, similar clusters in STS for both conditions, as well as bilateral superior parietal lobule and sPCS in self-generated imagery, were also identified using this perception searchlight.

After identifying brain clusters with robust orientation representations in the two imagery conditions, we next seek to identify clusters with significant representational differences between the two. We found significant lateralization of representational differences between conditions, with right sPCS demonstrating stronger orientation representations in self-generated imagery, and right extrastriate cortex demonstrating stronger orientation representations in cue-induced imagery (Figure 3C). To better illustrate the effects, we extracted multi-voxel activation patterns from the two regions of interest (ROIs) and generated reconstructions of imagined orientations of both conditions in each ROI. The representational strength of orientations in self-generated imagery was significantly higher than that in cue-induced imagery in right sPCS (*t*(19) = 3.13, *p* = 0.003). Notably, orientation reconstruction was significant in self-generated imagery (*t*(19) = 3.54, *p* = 0.001), but not in cue-induced imagery (*t*(19) = 0.69, *p* = 0.250). In addition, the representational strength of orientations in cue-induced imagery was higher than that in self-generated imagery in right extrastriate cortex (*t*(19) = 3.76, *p* = 0.001), with significant orientation representations in both conditions (*t*(19) = 6.32, *p* < 0.001 in cue-induced imagery, *t*(19) = 3.79, *p* = 0.001 in self-generated imagery). This posterior versus anterior differences in orientation representations resembled our findings in Experiment 1 which showed differential results in posterior and anterior electrodes.

If enhanced neural representations of orientations in sPCS supported the generation and maintenance of self-generated imagery, we would anticipate the representational strength of orientations in this region should be predictive of that in lower-level extrastriate cortex. Indeed, Pearson correlation analysis between the two revealed significant positive correlation in self-generated imagery (Figure S6B), *r* = 0.48, *p* = 0.034, but less so in cue-induced imagery (Figure S6A), *r* = 0.4, *p* = 0.077.

Lastly, as a control, we repeated the searchlight analysis on data from an earlier epoch of the trial (Figure 4, 6-9 s; 3 s after the retrocue). The representational strength of orientations in cue-induced imagery was significantly higher than that in self-generated imagery in bilateral V1 (*t*(19) = 3.76, *p* = 0.001) and right extrastriate cortex (*t*(19) = 4.29, *p* < 0.001; Figure 4C). These results were consistent with a previous fMRI study demonstrating successful decoding of retrieved stimulus-driven memories in early visual cortex following the onset of an associative cue ^5^. On the other hand, although there was no significant difference in right sPCS, there was a significant cluster in right sPCS in self-generated (Figure 4B) but not in cue-induced imagery (Figure 4A), suggesting that involvement of sPCS in self-generated imagery started early in the trial and became progressively larger into late memory delay.

**Figure 4.**
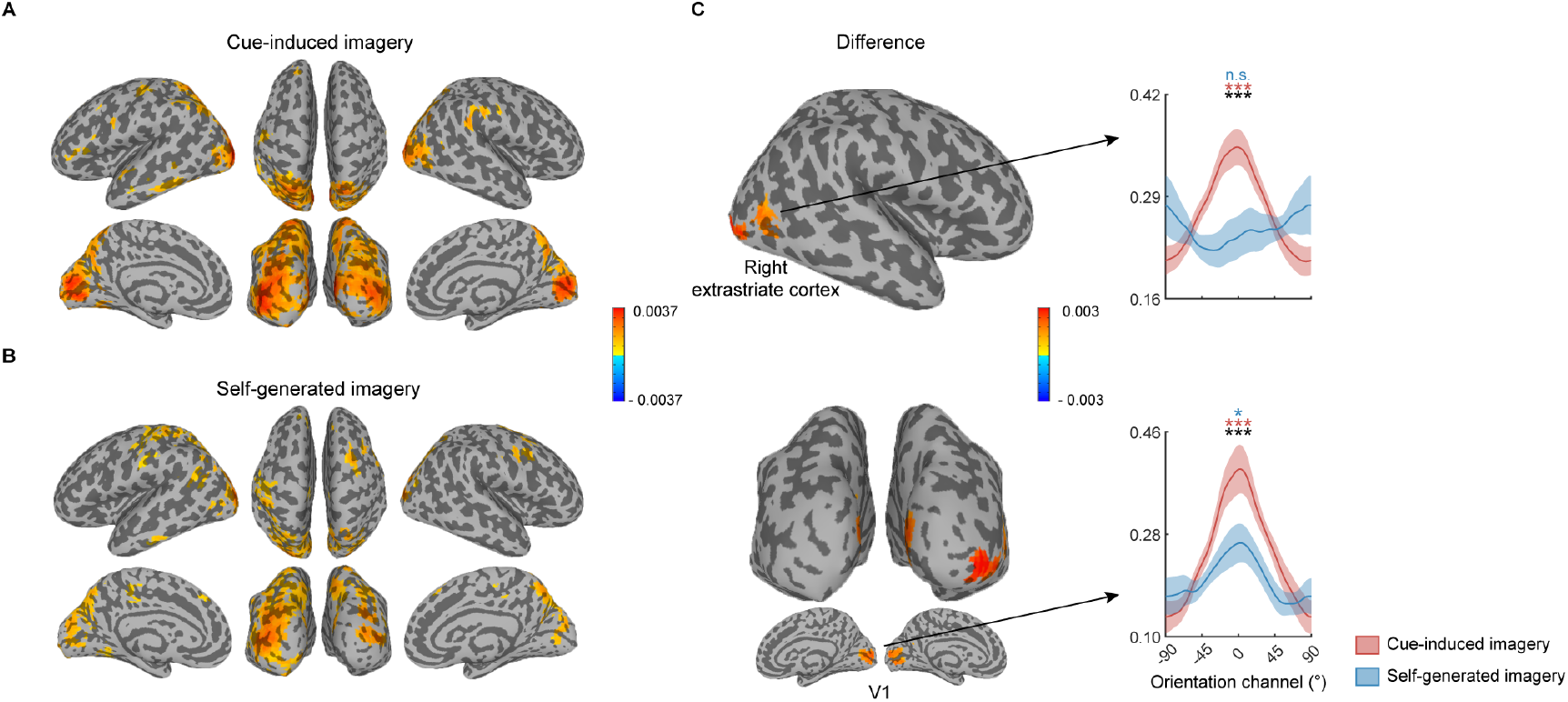
Whole-brain neural representations of imagery contents in middle delay. A. Searchlight parametric map of the strength of orientation representations in middle delay (6-9 s; 3 s after retrocue) in cue-induced imagery. Colors on the cortical surface denote brain regions with significant orientation representations, corrected using a cluster-based permutation method (*p* < 0.01). For demonstration purposes, clusters were thresholded at 50 voxels. B. Same as A, but with results from self-generated imagery. C. Difference map of orientation representations in A and B, with positive values denoting stronger orientation representations in cue-induced imagery. Orientation reconstructions obtained from the two significant clusters were shown, both clusters showed stronger orientation representations in cue-induced imagery: in right extrastriate cortex, only orientation representations in cue-induced imagery were significant: *t*(19) = 4.05, *p* < 0.001 in cue-induced imagery, *t*(19) = 0.61, *p* = 0.724 in self-generated imagery. In V1, orientation representations in both conditions were significant, *t*(19) = 4.15, *p* < 0.001 in cue-induced imagery, *t*(19) = 1.86, *p* = 0.039 in self-generated imagery. X axis represents distance from response orientations, with 0 representing the response orientation of each trial. Y axis represents reconstructed orientation channel responses in arbitrary units. Colored asterisks denote significance of cue-induced (red) and self-generated (blue) imagery, and black asterisk denotes significance of difference between conditions. n.s., not significant, *: *p* < 0.05, **: *p* < 0.01, ***: *p* < 0.001.

## Discussion

Even in the absence of any external stimulation, people can still self-generate contents in visual imagery. How would the neural underpinnings of self-generated imagery differ from those of classic cue-induced imagery? Here, we investigated (1) the temporal dynamics of self-generated imagery in an EEG experiment and (2) the spatial layouts of neural representations in self-generated imagery in an fMRI experiment, and contrasted the spatiotemporal dynamics of self-generated imagery with those of cue-induced imagery. Our results revealed an enhanced involvement of frontal cortex in generating and maintaining contents in self-generated imagery, as evidenced by enhanced neural representations of orientations in sustained potentials in central channels in EEG, and in sPCS of frontal cortex in fMRI. By contrast, cue-induced imagery was supported by enhanced neural representations of orientations in alpha-band activity in posterior channels in EEG, and in visual cortex in fMRI. Taken together, these results jointly support a reverse cortical hierarchy in representing imagery contents in self-generated versus externally-cued imagery.

Previous work on visual imagery has mostly utilized cue-induced paradigms, namely the to-be-imagined contents were guided by externally presented cues, either from long-term ^2-5^ or working memory ^6-9^. One advantage of using cue-induced imagery paradigms is that contents of imagery can be better controlled, compared to uncontrollable situations such as mind wandering. However, real-life imagery often requires imagery contents to be generated freely of external controls, yet the neural mechanisms of self-generated imagery have remained largely unexplored due to limitations in experimental paradigms. Although there have been several recent attempts to tackle on a related problem ^26,27^, it has remained unclear how contents of self-generated imagery were generated and how self-generated imagery differed from classic cue-induced imagery. To balance the needs for well-controlled experimental settings as well as for self-generating imagery contents, here we designed a novel experimental paradigm to investigate the neural mechanism of self-generated imagery: participants decided freely what to imagine on each trial, with the constraint that imagery contents were limited to a set of seven pre-learned line orientations. Combining the new behavioral paradigm with multivariate inverted encoding models allowed us to examine the neural representations of self-generated imagery contents (orientations in our case), and how these representations differed from those of cue-induced imagery, beyond univariate activation changes between conditions. The present study revealed several distinctive features of self-generated imagery: behaviorally, the vividness of self-generated imagery was significantly reduced compared to that of cue-induced imagery; neurally, self-generated imagery shared representational codes with perception as well as with cue-induced imagery in early visual cortex; more interestingly, converging evidence from EEG and fMRI suggested enhanced orientation representations in frontal cortex in self-generated compared to cue-induced imagery. We interpreted these representational differences as reflecting a reverse cortical hierarchy in representing imagery contents that were generated either via internal drives or external cues. Below we discuss our findings in EEG and fMRI in more details:

According to the reverse visual hierarchy model, imagery signals are initiated in the more anterior part of cortex such as frontal cortex, and the signals trigger a cascade of neural processing along the anterior-to-posterior cortical hierarchy ^1,17^. Because the initiation signals in self-generated and cue-induced imagery derived from completely different origins, we hypothesized that anterior cortex would act differently in self-generated and cue-induced imagery. Our results from both EEG and fMRI supported this notion: in EEG, we found that imagery contents were decodable in sustained potentials in central but not in posterior electrodes in both conditions, and more importantly, orientation representations emerged earlier in time in self-generated imagery, and remained stronger than those in cue-induced imagery in central electrodes. Due to the poor spatial resolution of EEG signals, we next turned to fMRI for the neural loci of such representational differences. Consistent with the EEG findings, we observed that right sPCS of frontal cortex maintained robust orientation representations of self-generated imagery but not cue-induced imagery, in both middle and late delay periods. Together, these results indicated anterior cortex, especially right sPCS in the current study, might serve as the critical neural locus that initiates and maintains contents in self-generated imagery.

The results of sPCS in the current study are broadly in line with previous work implicating a role of sPCS in visual working memory ^20-22^. Our results extended this finding to the imagery domain, and more specifically, we demonstrated that sPCS contributed to self-generated imagery in a way that was specific to imagined stimuli. Recent debates in the field of working memory have argued about the specific role of higher-order frontal cortex in working memory maintenance ^28,29^, partially due to the fact that stimulus-specific representations observed in frontal cortex during working memory were substantially more variable compared to low-level visual cortex. Our finding added new insights into this line of research, by demonstrating that stimulus-specific representations in frontal cortex were enhanced when the level of “internality” increased as in self-generated imagery. In other words, our work clearly indicated the origin of stimulus-specific representations in sPCS was internal rather than external. Moreover, we demonstrated significant functional coupling between stimulus-specific representations in sPCS and those in EVC in self-generated imagery, in support of the view that sPCS exerts top-down control over lower-level visual cortex ^30,31^. Intriguingly, in another recent work from our lab (unpublished), we have identified a similar reverse hierarchy between EVC and IPS, for cue-induced imagery as compared to perception. The fact that the top node of the reverse hierarchy moved more anteriorly from IPS to sPCS, when imagery contents became more “internally-generated”, possibly implied a flexible reverse hierarchy that depends on the “internality” of the specific cognitive process. Last but not least, it should be noted that the function of sPCS can be better understood when taking into account the type of imagined stimuli used in the current study. sPCS shows robust stimulus representations for spatial or space-related stimuli such as locations and orientations ^20-22^, but less so for non-spatial stimuli such as colors ^21,32^. Whether there remains a more domain-general, stimulus-nonspecific brain region in self-generated imagery requires further future work to elaborate on.

Turning to lower-level visual cortex, imagery and perception have been shown to share neural codes in early visual cortex ^8,33^. Relatedly, a recent EEG experiment found imagery and perception shared neural representations in alpha-band oscillatory activity in posterior electrodes ^4^. In our EEG experiment, although we failed to find orientation representations from sustained potentials in posterior electrodes, we found imagery contents were indeed represented in alpha-band oscillatory activity in posterior electrodes. Interestingly, orientation representations in alpha-band activity demonstrated a reversed pattern from those in sustained potentials, and orientation representations were stronger in cue-induced imagery rather than in self-generated imagery. Our fMRI searchlight also demonstrated an analogous pattern that first emerged in V1 and moved onto extrastriate cortex in late delay, and orientation representations of both self-generated and cue-induced imagery shared common neural codes with perception in visual cortex. Moreover, the emergence of orientation representations in posterior electrodes was much later in time, compared to those in central electrodes in sustained potentials. Given that alpha-band oscillations carry feedback information ^34-36^, it was likely that the orientation representations in alpha-band received feedback modulations from higher-order areas, possibly frontal cortex.

How should we interpret the reversed patterns of results in alpha-band activity in EEG as well as in visual cortex in fMRI? We noticed that this neural result echoed the behavioral difference in vividness, that is, imagery experience was reported to be more vivid in cue-induced imagery than in self-generated imagery. One explanation for the reduced representational strength in self-generated imagery would be attenuated sensory processing for self-generated imagery, similar as self-generated sensations that felt less salient than externally generated sensations ^37-39^, and as other higher-level self-generated cognitive processes such as motor imagery, inner speech, and numerosity estimation ^40-42^. Attenuation in self-generated imagery can be accommodated within the internal feedforward model framework ^43^. This model proposes any action people take is followed by a corollary discharge, which is used to predict sensory consequences of the action. When the prediction matches the actual sensory feedback, the sensory consequences are attenuated. In self-generated imagery, the prediction of imagery was always in line with self-generated contents, thus leading to weaker neural representations in sensory cortex. Consequently, the reduction in subjective experience of imagery might derive from this representational attenuation.

Alternatively, these results could be accounted for by the sensorimotor recruitment hypothesis, which proposes that visual cortex is engaged in both perceiving external stimuli and maintaining mental images ^44^, with shared representations between perception and working memory ^45^, between long-term memory and perception ^5^, and between long-term memory and working memory ^46^. In these studies, significant neural representations of contents in long-term memory in early visual cortex can be explained by a neural reinstatement of the to-be-retrieved information from long-term memory, with the hippocampus possibly acting as the source of the modulatory signals ^5^. Because both of the imagery tasks in our current study engaged retrieval from long-term memory, it was possible that the associative cue in cue-induced imagery facilitated memory retrieval and resulted in a stronger reinstatement of imagined contents. It should be noted that the sensory attenuation and sensorimotor recruitment accounts are not mutually exclusive, and might simultaneously contribute to the current results.

It is noteworthy that in our EEG experiment, distinct result patterns were observed in sustained potentials and oscillatory activity: first, orientation representations were observed in central electrodes in sustained potentials, and in posterior electrodes in alpha-band activity; second, the differences in representational strength between self-generated and cue-induced imagery were reversed, with self-generated imagery demonstrating better orientation representation in central electrodes in sustained potentials, and the opposite was true in posterior electrodes in alpha-band activity in cue-induced imagery. While these results might seem difficult to explain at first glance, we would like to point out that several recent studies have also reported distinct cognitive processes carried by sustained potentials and oscillatory activity. For example, one study has found that sustained potentials encoded contents in working memory, whereas alpha-band activity mainly encoded spatial attention ^24^. However, in a different memory paradigm, unattended items in working memory could be decoded from alpha-band activity but not from sustained potentials ^25,47^. We think these results would be difficult to reconcile, without a systematic examination on the effects of the two types of EEG activity along with careful experimental designs; yet, we speculated that narrowing down the focus to alpha-band activity by applying wavelet transformation to sustained potentials might filter out orientation-irrelevant signals and noise in other frequency bands, thereby increasing the signal-noise ratio (SNR) of orientation representations in posterior electrodes. By contrast, imagery-relevant signals in central electrodes might rely primarily on slow-wave cortical dynamics ^48^, because the pattern of enhanced self-generated representations was not observed in any single frequency band of oscillatory activity in central electrodes. It would be interesting for future studies to examine whether the observed functional differences of frontocentral sustained potentials and posterior alpha-band activity would generalize to other cognitive processes.

We have discussed several distinct features of self-generated imagery by comparing it with cue-induced imagery; yet, there remain a few other interesting observations from the current study that require further work to look into. For instance, we noticed that the majority of differences between cue-induced and self-generated imagery was cortically right lateralized. Moreover, in terms of the involvement of frontal cortex, there was a hint that self-generated imagery was right lateralized, and cue-induced imagery was left lateralized. Whether this differential patterns in cortical lateralization speaks to difference between different types of imagery remains to be further investigated ^49^. In addition, although we removed any potential response bias from the model training stage to avoid overfitting, it would be interesting to investigate whether the two types of imagery are influenced by different cognitive factors and thereby resulting in differential response bias patterns, such as the oblique and attractor biases typically observed in working memory ^50,51^.

In conclusion, using both EEG and fMRI, we revealed distinctive spatiotemporal neural dynamics underlying the neural basis of self-generated imagery: compared to cue-induced imagery, self-generated imagery was supported by an enhanced involvement of frontal cortex, as indexed by better imagery representations in sustained potentials in central channels of EEG and in sPCS of frontal cortex in fMRI. The enhancement in frontal representations was accompanied by a decrease in orientation representations in visual cortex, which might reflect the attenuated subjective experience in vividness at the behavioral level. Research on self-generated imagery may have abundant potential uses. People who suffer from schizophrenia might either have delusions that have no basis in reality or generate hallucinations whose contents do not actually exist. Our results provide new insights into the neural mechanisms of visual imagery, and may open up a new avenue for both experimental and clinical research on imagery.

## Methods

### Participants

A total of forty-nine volunteers participated in the study, two of whom participated in both experiments. All participants had normal or corrected-to-normal vision, reported having no psychiatric or neurological disorders, provided written informed consent, and reported normal visual imagery ability assessed by the Vividness of Visual Imagery Questionnaire (VVIQ) ^52^. All participants were recruited at Shanghai Institutes for Biological Sciences, Chinese Academy of Sciences, and were monetarily compensated for their participation. The study was approved by the ethical committee of Center for Excellence in Brain Science and Intelligence Technology, Chinese Academy of Sciences (CEBSIT-2020028).

Twenty-eight volunteers participated in Experiment 1 (EEG experiment). Four participants were excluded: two participants had insufficient data due to technical issues, one participant failed to follow instructions, and one participant dropped out from the experiment, leaving twenty-four participants in the final sample for Experiment 1 (13 females, 11 males; mean age = 24.1, SD = 2.3). Twenty-three volunteers took part in Experiment 2 (fMRI experiment), all were eligible for MRI scans. Three participants were excluded: two participants had insufficient data due to technical issues, one participant failed to follow instructions, leaving twenty participants in the final sample for Experiment 2 (11 females, 9 males; mean age = 23.6, SD = 2.3). We did not estimate sample sizes for Experiment 1 or 2 a priori, but the sample size used in both experiments were comparable to those in previous studies with similar approaches.

### Stimuli & Apparatus

Two sets of non-semantic kaleidoscope images were used, each consisted of seven images. All of them were generated by Python2 codes used in a previous study ^53^. Each kaleidoscope was created by overlaying three transformed hexagons. Each hexagon had a unique color and was transformed by four rounds of side deflection at a random direction. The RGB values of colors were [(220,20,60), (70,130,180), (255,140,0)] in one set and [(50,205,50), (139,0,139), (205,155,29)] in the other set.

In Experiment 1, stimuli were presented with MATLAB (R2018b, The MathWorks) and Psychtoolbox 3 extensions ^54,55^. They were displayed on a 48×27 cm HIKVISION LCD screen with a 60 Hz refresh rate and a 1920 × 1080 resolution. The viewing distance was 62 cm. While performing the task, participants’ head position was stabilized by a chin rest. Responses were recorded with a keyboard and a mouse.

In Experiment 2, All stimuli were presented using MATLAB (R2012b, The MathWorks) and Psychtoolbox 3 extensions on an SINORAD LCD projector (1280 × 1024 resolution; 60 Hz refresh rate). Participants viewed stimuli through a coil-mounted mirror in the scanner at a viewing distance of 90.5 cm. Responses were made via two SINORAD two-key button boxes.

### Experimental paradigm and procedure

#### Overview

The purpose of the current study was to unveil the spatiotemporal neural dynamics of self-generated imagery, by contrasting with those of cue-induced imagery. In both conditions, we presented kaleidoscope images instead of the actual to-be-imagined stimuli, in order to minimize the influence of stimulus-driven activity in neural signals. In cue-induced imagery, imagery content was determined by an external cue. Seven kaleidoscope images were used, each associated with a specific line orientation. Participants were required to imagine a line with the orientation indicated by the kaleidoscope image. To further eliminate stimulus-driven activity from the kaleidoscope images, we adopted a retrocue imagery paradigm. On each trial, participants were presented with two kaleidoscope images followed by a retrocue. The retrocue indicated the specific kaleidoscope image with which the associated orientation should be imagined. In self-generated imagery, the imagery content was determined by participants volitionally. Participants needed to generate their imagery content on their own by freely choosing one from the seven learned orientations on each trial. To match the trial time course of the cue-induced imagery, seven different kaleidoscope images and a retrocue design were also used, but the kaleidoscopes were not associated with orientations. The experimental paradigm was depicted in Figure 1. The specific set of kaleidoscope images used for each condition was counterbalanced across participants. The specific association between each kaleidoscope image and each orientation was also randomized across participants.

Another goal of the current study was to obtain trialwise objective and subjective measures of imagery. At the end of each trial, participants were required to report their imagery content (orientation) and subjective vividness. In Experiment 2, due to time limitations, we used 1-4 point of vividness rating for assessing subjective vividness as used in previous studies ^56^. However, the standard measurement of vividness such as 1-4 point rating conflated lots of different factors of subjective experience, such as subjective specificity and subjective intensity ^57^. As such, to uncover specific subjective experience in different dimensions, in Experiment 1 the vividness was decomposed into two different dimensions: precision and intensity. Precision represented the confidence in the precision of orientation report, whereas intensity indicated the subjective strength of imagery content.

#### Behavioral learning session

Prior to the main experimental session, participants first learned the associations between seven kaleidoscope images and seven specific orientations (spanning the entire orientation space and were equally distant: 15°, 40.71°, 66.43°, 92.14°, 117.86°, 143.57°, 169.29°). On each trial, participants passively viewed one kaleidoscope image followed by its associated line orientation. The kaleidoscope image was presented for 0.9 s and then the line was shown for 0.5 s, with an inter-stimulus-interval (ISI) of 0.2 s in between, followed by an inter-trial-interval (ITI) of 1.2 s. Each learning block consisted of all seven association pairs presented in a randomized order. At the end of each block, participants could decide whether to perform a test on their learned associations or to continue with learning. Each trial started with a fixation period of 0.5 s, and then a kaleidoscope image was presented for 0.4 s. After a 2-s delay, participants were required to report the corresponding orientation on an orientation wheel as precisely as possible in 2.5 s. A feedback message would be presented for 0.3 s, indicating whether the response was accurate (error < 5°) or inaccurate (error >= 5°). After an interval of 0.2 s, the line with the correct orientation would be shown for 0.5 s to consolidate memory. ITI varied in 0.8-1.3 s. The test phase consisted of 28 trials. The radius of the line in both learning and testing phases varied between 3.2-4.8° on a trial-by-trial basis. Participants underwent the aforementioned procedure iteratively until the mean absolute error during test fell below 10°.

#### Experiment 1

In Experiment 1 (the EEG experiment), each trial started with the successive presentation of two kaleidoscope images at the center of the screen (3.65°×3.65° in size), each for 0.35 s with an ISI of 0.2 s. After 0.2 s, a retrocue followed for 0.2 s indicating which of the two stimuli should be used for imagery. If images had no associations (self-generated imagery), participants needed to freely choose one from seven learned orientations and imagine a line with the chosen orientation at the center of the screen; if images were associated with orientations (cue-induced imagery), participants needed to imagine the line at the orientation associated with the cued kaleidoscope image. After a delay period of 2 s, during which participants needed to keep actively imagining the orientation, participants were required to report the orientation, precision and intensity of their imagined line on an orientation wheel in 5.5 s. The orientation wheel consisted of a circle with a radius of 5°, a needle crossing the fixation point with the same radius, and a bowtie-shaped wedge centered on the needle. The orientation of the needle represented the orientation of the imagined line, which was adjusted by changing the position of the mouse cursor. The angle of wedge represented precision, which was adjusted by changing the distance of cursor to the fixation. The color of the wedge indicated intensity, which was adjusted by two buttons (increase or decrease) on a keyboard. The initial values of orientation, angle and color were randomly chosen and participants could move the cursor and press keyboard to report three variates simultaneously. Only when both operations of mouse and keyboard were finished would the trial end. ITI varied in 0.8-1.3 s. The cued kaleidoscope, cued order (first versus second), and condition were fully counterbalanced across blocks.

In each block, there were three catch trials to keep participants’ attention on kaleidoscope images before delay. The catch trial had the same time course as the main task trial, except that it required participants to choose the cued kaleidoscope from two probe images after retrocue in 3 s. catch trial ITIs were fixed at 1.05 s. At the end of each block, participants received feedback on main task performance and catch trial performance. Taking account of catch trials, there were 45 trials per block. All participants needed to complete 20 blocks in Experiment 1. In total, participants performed 420 trials per condition. Seven participants performed the task without catch trials.

#### Experiment 2

##### Imagery task

The procedure of Experiment 2 was similar to that of Experiment 1, except that the timing of events and responses were adjusted for fMRI. Participants were shown two kaleidoscope images (3.26°×3.26° in size) successively, each in a 0.8-s stimulus window with a 0.4-s ISI. Then a retrocue was presented for 0.6 s. During the delay period, participants imagined a line for 9 s. During the response period, participants needed to rotate the needle of orientation wheel (radius = 3.7°) to match the orientation of imagined line within 3.75 s and then rated their experienced vividness on a scale from 1 to 4 points in 1.25 s, where 1 represented lowest vividness and 4 represented highest vividness. The ITI varied in 2.5 s, 4 s, 5.5 s and 7 s. Task performance and the number of missing vividness reports were provided at the end of each block as feedback. There were 16 main task trials and one catch trial per block. The response time for catch trials was 3 s, and catch trial ITIs were fixed at 4.5 s. Each participant completed 14 blocks in total, resulting in 112 trials per condition. Three participants performed the task without catch trials.

##### Perception task

Because no physical orientations were present throughout the imagery task, in order to obtain participants’ neural responses to ground-truth, sensory orientations, participants completed three additional blocks of perception task following the imagery task. On each trial, an oriented line whose orientation was randomly chosen from seven specific orientations flickered at the center of the screen for 4.5 s at a frequency of about 1.8 Hz. The radius of the line randomly varied between 3.2-4.8° on a trial-by-trial basis. The ITI varied in 3 s, 4.5 s and 6 s. Participants were instructed to fixate at the white fixation point and press a corresponding button whenever the fixation point turned green. Each perception block consisted of 30 trials, and participants completed 90 trials in total.

##### EEG recording and preprocessing

EEG data were acquired using a Brain Products ActiCHamp recording system and BrainVision Recorder (Brain Products GmbH, Gilching, Germany). Scalp voltage was obtained from a broad set of 59 electrodes at 1000 Hz (FCz as reference). Vertical and horizontal EOG were recorded from 2 electrodes located ∼2 cm above and below the right eye, and from 2 electrodes ∼1.5 cm lateral to the external canthi, respectively. Electrode impedance was kept below 30 kΩ.

Preprocessing analyses were performed in MATLAB (R2021a, The MathWorks) using EEGLAB Toolbox ^58^. The raw EEG signals were resampled at 250 Hz. Then the data were band-pass filtered between 0.01 and 45 Hz. Epochs were segmented from -0.5 s to +3.6 s relative to the onset of the first stimulus. The signals were baseline corrected from -0.2 s to 0 s. The epoched data were visually inspected and those containing large muscle, cardiac and respiratory artifacts (except for eye blinks) or extreme voltage offsets were manually removed. Independent component analysis (ICA) was then performed using EEGLAB’s binica algorithm for each subject to identify and remove components that were associated with eye blinks ^59^ and eye movements ^60^. Data after ICA were treated as the preprocessed voltage data. To estimate oscillatory power across time and frequencies, the voltage data from each channel and trial were convolved with a family of complex Morlet wavelets spanning 3–45 Hz in 1 Hz steps with wavelet cycles increasing linearly between 3 and 10 cycles as a function of frequency. The power was calculated as the percent change of squared absolute value in the resulting complex time series relative to the baseline between −0.2 s and 0 s.

##### fMRI acquisition and fMRI data preprocessing

MRI data were recorded using a Siemens Tim Trio 3.0 T scanner (Erlangen, Germany) with a standard 32-channel phased-array head coil at the Center for Excellence in Brain Science and Intelligence Technology, Chinese Academy of Sciences. Functional images were acquired with a gradient echo echoplanar pulse sequence with a multiband acceleration factor of 2 (TR/TE = 1500/30 ms; flip angle = 60°; matrix = 74 × 74; 46 slices; voxel size = 3 mm isotropic). T1-weighted anatomic images were collected using the Magnetization Prepared Rapid Acquisition Gradient Echo (MPRAGE) pulse sequence (TR/TE = 2300/2.98 ms; flip angle = 9°; matrix = 256 × 256; 192 slices; voxel size = 1 mm isotropic).

Preprocessing of MRI data was performed using AFNI ^61^. The first five volumes of each functional run were removed. The EPI data were then registered to the last volume of each scan session and then to the T1 volume of the same session. Six nuisance regressors were included in GLMs to account for head motion artifacts in six different directions. The data were then motion corrected, detrended (linear, quadratic, cubic), and z-score normalized within each run.

##### Quantification and statistical analyses

###### Behavioral data analyses

In both Experiments 1 and 2, the behavioral performance of imagery could be quantified by errors of the responses relative to the ground-truth orientations. In cue-induced imagery, error was calculated as the circular distance between the cued and response orientations, which we referred as the absolute error. In self-generated imagery, because there were no sample orientations in self-generated imagery, errors in this condition were quantified by calculating the circular distance of response orientations to all the seven learned orientations, and choosing the smallest error among all as the relative error. As a comparison, relative errors were also computed for cue-induced imagery. For vividness measurements, in Experiment 1, precision was quantified by the angle of response wedge (ranging from 2° to 180°), of which the smaller angle represented smaller uncertainty of the orientation of imagined line. Intensity was quantified by the grayscale value of response wedge (ranging from 0 to 0.5; 0 = black, 0.5 = background color). Smaller values represented a more intensive imagery experience. Thus, the smaller the values of precision and intensity, the more vivid the subjective experience. In Experiment 2, the vividness rating score represented the level of vividness, the larger the vividness score, the more vivid the subjective experience.

For each participant, means of error, precision, intensity or vividness rating in each condition were calculated. We conducted two-tailed paired t tests to test the significance of the mean difference between conditions. Pearson correlations were performed between precision, intensity and error to assess their potential correlational relationships.

We took several approaches to assess whether there existed any systematic biases in participants’ responses. First, we examined the uniformity of response distributions in two conditions. All responses were binned into seven bins, with each of the seven learned orientations as bin centers. Differences in distributions were statistically assessed using ***χ***^2^ tests. Second, we examined whether the initial orientation had a systematic influence on response, by calculating the circular distance between the initial orientation and final response orientation. Differences between bins were statistically assessed using two-tailed paired t tests.

#### Inverted encoding model (IEM)

##### Overview

All IEM analyses were performed in MATLAB using custom codes. The inverted encoding model assumed that the signals in each unit (e.g., voltage or power in each electrode in EEG, or BOLD signal in each voxel in fMRI) reflected the weighted sum of a small number of hypothesized feature tuning channels (i.e., neuronal populations), each tuned for a different feature (i.e., orientation in the current study). In our experiments, the number of hypothesized orientation tuning channels was set to five (36° apart, equally spaced). We modeled the response profile of each channel to a specific orientation θ as a half sinusoid raised to the 8th power (FWHM = 0.82 rad):

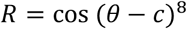

where c was the center of the channel. Since there were no correct targets in self-generated imagery, we took response orientations (round to the nearest integer) in both conditions as θ to obtain the idealized responses from basis functions, which meant the θ was possible in the 1-180° orientation space.

The IEM analysis proceeded in two stages, encoding (training) and decoding (test). We partitioned our data into independent sets of training data and test data. In the encoding stage, the hypothesized channel responses (C_1_, k × n, k: the number of channels; n: the number of trials) were projected to actual measured signals in training dataset (B_1_, m × n, m: the number of units) according to an unknown weight matrix (W, m × k), which could be described a general linear model of the following form:

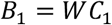

The weight matrix (Ŵ) was obtained via least-squares estimation as follows:

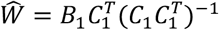

In decoding stage, the model was inverted to transform the independent test dataset (B_2_, m × t, t: the number of trials) into estimated channel responses (C_2_, k × t) by the obtained weight matrix:

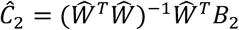

Following the IEM analysis in previous studies ^27,62^, the channel centers were not fixed but shifted from 0°, 36°, 72°, 108°, 144° to 35°, 71°, 107°, 143°, 179° in 1° step for 36 iterations. We conducted the above analysis in each iteration, such that all 180 orientations from 1° to 180° served as channel centers. All of estimated channel responses from all iterations were combined to create responses of 180 orientation channels. The result, for any given orientation, can be considered a reconstruction of the model’s estimate of the neural representation of that orientation. This procedure ensured that our reconstructions were not biased by any specific channel centers. The reconstruction of channel responses was shifted to a common center (90° on x axis).

To characterize the strength of reconstructions, we folded the channel responses on both sides of the common center, averaged them, fitted with linear regression, and then took the resulting slope of linear regression as an index of the strength of reconstructions.

##### IEM procedure with all and balanced data

To reveal orientation-specific neural representations of imagery in both conditions, we used participants’ response on each trial as the target label, and used data combined from both conditions for training and testing IEMs. This mixed IEM was supposed to provide an unbiased way of making comparisons between conditions ^63^. To achieve this, we used a k-fold cross-validation procedure. For each participant, all data from both conditions were divided into four folds. In each iteration, all but one folds served as the training data, and the left-out fold served as the testing data. The procedure iterated until all folds had served as training and testing data, and results from all iterations were averaged, for each condition separately.

However, one potential drawback with the approach was that participants’ responses were often unbalanced across different response bins. This imbalance in trial number between response bins might result in overfitting of IEM, such that orientation reconstructions from IEM might be overestimated. To avoid this, we balanced trials in a way that the trial number in each of the seven bins would be made equal. To be specific, we randomly drew a certain number of trials from each bin, and the number of trials drawn was determined by the bin with the smallest number of trials among the seven bins. This initial step would result in matched numbers of trials across bins, but not necessarily between conditions. To further balance trials between conditions, we rebalanced the trials by randomly removing one trial from a certain bin of the condition with more trials; and in the meantime, including one trial from the same bin of the condition with fewer trials, this step was iterated until the difference of trial numbers between conditions was below two. To make full use of all data, the balancing and cross-validation procedures were repeated for 50 times, and the results were averaged across repetitions.

##### IEM analyses with EEG

In EEG experiment, 59 electrodes were divided into 3 subsets of electrodes (frontal electrodes: FP1, FP2, AFz3, AFz4, AFz7, AFz8, Fz1, Fz2, Fz3, Fz4, Fz5, Fz6, Fz7, Fz8; central electrodes: FC1, FC2, FC3, FC4, FC5, FC6, FT7, FT8, Cz1, Cz2, Cz3, Cz4, Cz5, Cz6, T7, T8, CPz1, CPz2, CPz3, CPz4, CPz5, CPz6, TP7, TP8; posterior electrodes: Pz1, Pz2, Pz3, Pz4, Pz5, Pz6, Pz7, Pz8, POz3, POz4, POz7, POz8, Oz1, Oz2). We applied IEM to voltage signals from all electrodes, frontal electrodes, central electrodes and posterior electrodes separately. After time-frequency decomposition of voltage data, the obtained power signals were averaged within alpha-band (8-12 Hz). Similarly, we performed IEM analyses on alpha-band power data, separately in global electrodes and local subsets. For both voltage and power data, after balancing trials, IEM analysis was performed at each time point with a sliding window of 3 time points to obtain time-resolved orientation reconstructions. For IEM analyses on all frequencies, a similar procedure was conducted on power data of every single frequency ranging from 3 to 45 Hz.

##### IEM analyses with fMRI (Searchlight of IEM)

In Experiment 2, the IEM was combined with a roving ‘‘searchlight’’ procedure ^20,64^, which allowed us to reconstruct and quantify representations of imagined orientations across the entire brain. For each participant, their data in the original space were warped to the MNI template ^65^. We used the “sphere_searchlight” class in PyMVPA toolbox ^66^ to perform the searchlight analysis. We defined a spherical searchlight (radius = 9 mm) centered on each voxel of the whole-brain gray matter mask. Considering a typical hemodynamic response lag of 4-6 s, we extracted and averaged the BOLD responses in each voxel over a time period spanning 6-9 s (middle delay) and another spanning 9-12 s (late delay) following the onset of stimulus, and performed IEM searchlight within each time period. IEM analysis was performed using the data with all trials and balanced trials to calculated the slope maps separately. Results from whole-brain searchlight were displayed on the cortical surface reconstructed with FreeSurfer ^67,68^ and visualized with SUMA in AFNI.

Because no physical lines were present during the imagery task, to compare the neural representation of imagery with that of perception, we performed a cross-task generalization IEM analysis, by training the IEM on perception data, and testing the IEM on imagery data. We extracted and averaged the responses in each voxel over a time period spanning 4.5-7.5 s of each trial of the perception task to train the IEM searchlight, and tested the model on the late delay data (9-12 s) of the imagery task. We did not balance trials for this analysis because samples in our perception task were already balanced.

Note that the total trial number in each condition was smaller in Experiment 2 than in Experiment 1 due to prolonged trial length in fMRI studies. To avoid false positive results from an insufficient number of trials, we conducted the searchlight with trial balancing and without trial balancing (i.e., using all trials), and took the intersection of these two statistical parametric maps as the final result of the searchlight analyses in Experiment 2.

#### Cluster-based multiple comparisons correction

We used cluster-based permutation to correct for multiple comparisons across time points (in EEG) and voxels (in fMRI). The overarching principle for cluster-based permutation is depicted as below: all to-be-corrected test statistics were clustered in connected sets on the basis of temporal, spatial, or spatiotemporal adjacency to form contiguous clusters. Cluster-level test statistics were calculated by taking the sum of the test statistics within every cluster. The test statistics were then permuted, and the cluster-level test statistic of the largest cluster was taken from the permuted data. This procedure was repeated 10000 times to create a null distribution of test statistics. The proportion of each cluster-level test statistic in true data being smaller than the cluster-level null distribution was calculated as the p-value of the corresponding cluster.

For analyses on voltage and alpha-band data of EEG, the IEM slopes from 24 participants in each condition and their paired difference between conditions were compared against zero using paired or one-sample t-test to obtain the one-tailed significance (α = 0.05) at each time point. To obtain the null distribution for multiple comparisons correction, the slopes at each time point were randomly multiplied by either 1 or -1 independently, and then performed one-tailed t-test across participants. This procedure was repeated 10000 times, creating a null distribution of the t statistics. The contiguous t statistics clusters of true data and the null distribution underwent multiple comparisons correction to threshold (α = 0.01) significant time points. For analyses on all frequency data of EEG, a similar procedure was conducted on 2D time-frequency clusters.

For fMRI data, t-tests for the whole-brain slope map in each condition and their difference against zero were performed using the AFNI function “3dttest++”. To reduce the computational load in permutation, we performed a two-stage procedure ^69^. The randomized sign-flip procedure described above was first repeated 100 times for each participant. In a second step, the randomized samples were bootstrapped from each participant and then performed one-tailed t-test (α = 0.05) across participants to obtain a map of t statistics. The second step was repeated 10000 times to create the null distribution. The contiguous t statistics clusters of true slope maps and the null distribution underwent multiple comparisons correction and the obtained *p*-values were further corrected using Family-wise Error Rate (FWE) method using a threshold of α = 0.01.

#### Classification using Support vector machine

In order to examine whether our main results would hold with a different approach for revealing orientation representations, we decoded imagery content using multi-class classification with a linear support vector machine (SVM) approach. We first labeled the responses according to which of the seven bins the responses belonged to. Then the trial number in each bin were balanced using the method mentioned above and a k-fold (k = 4) cross-validation procedure was applied to the data with balanced trials. Like the IEM analysis with balanced trials, the SVM decoding was also repeated 50 times and the results from each iteration were averaged. We used the “fitcecoc()” function with standard linear SVM classifier as the learner to decode the EEG data.

## Supplementary Figures

**Figure S1.**
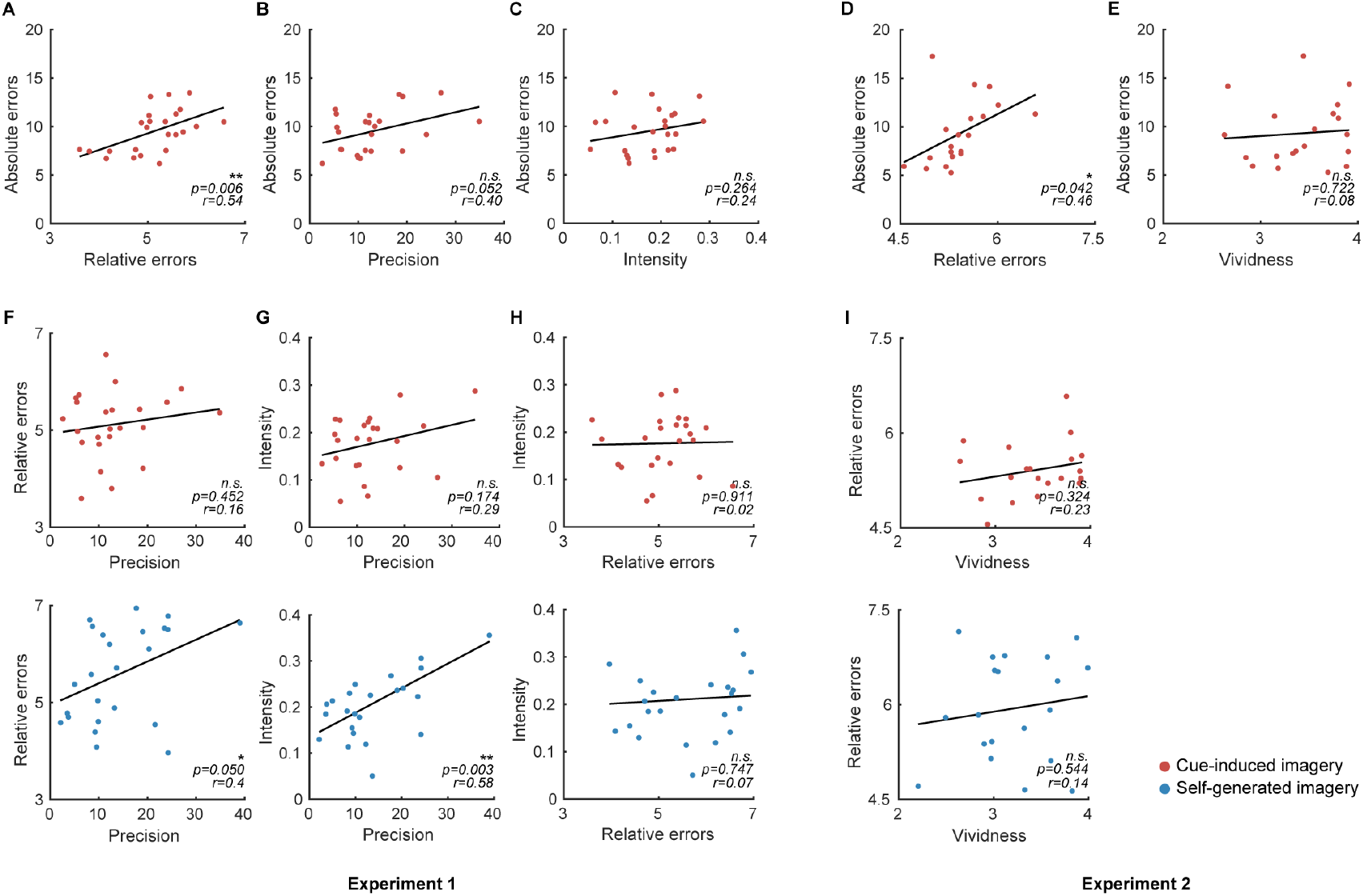
Correlations between behavioral measurements across participants. A. Relative errors positively correlated with absolute errors in cue-induced imagery in Experiment 1. Each dot represented individual participant. Relative errors (x axis) and absolute errors (y axis) were averaged across trials for each participant. Black lines represented the best linear fit. B. Similar as A, but with correlations between precision and absolute errors in Experiment 1. C. Similar as A, but with correlations between intensity and absolute errors in Experiment 1. D. Same as A, but with results from Experiment 2. E. Similar as A, but with correlations between vividness and absolute errors in Experiment 2. F. Similar as A, but with correlations between precision and relative errors in Experiment 1. G. Similar as A, but with correlations between precision and intensity in Experiment 1. H. Similar as A, but with correlations between relative errors and intensity in Experiment 1. I. Similar as A, but with correlations between vividness and relative errors in Experiment 2. Red and blue dots represent cue-induced and self-generated imagery, respectively. Asterisks denote significance of correlations, n.s., not significant, *: *p* < 0.05, **: *p* < 0.01, ***: *p* < 0.001.

**Figure S2.**
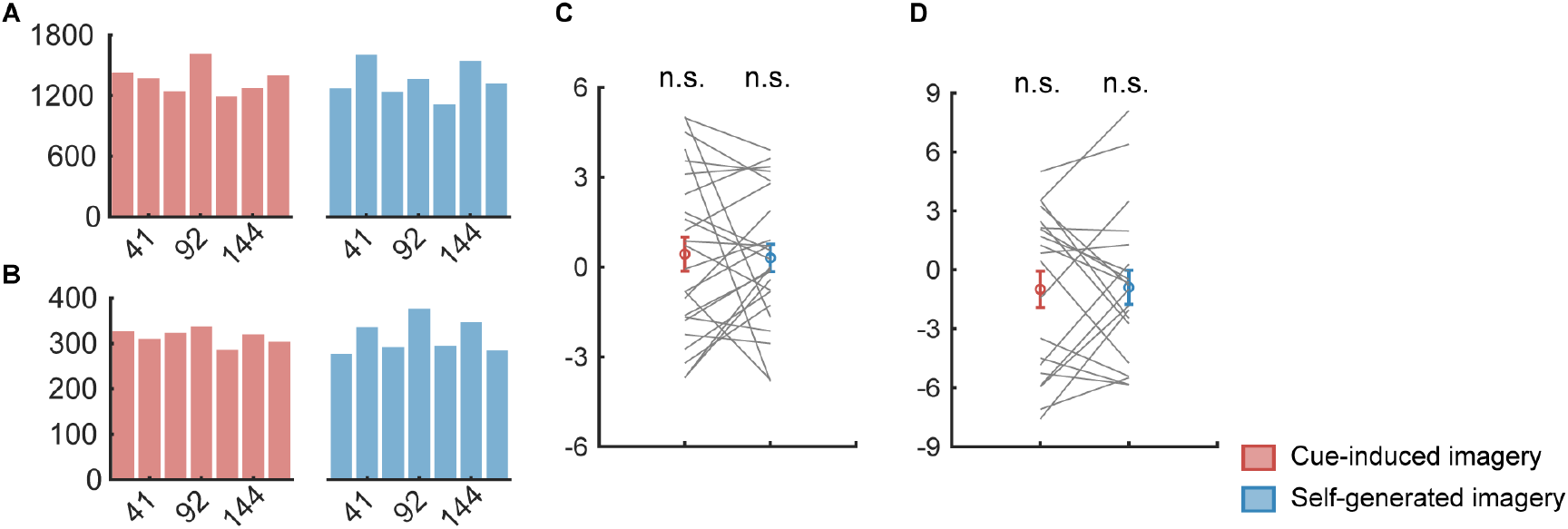
Evaluation of response biases. A. Histograms of response distributions, pooled from all participants in Experiment 1. x axis represents orientation bins, centered on the seven learned orientations, y axis represents frequency. Red and blue bars represent cue-induced and self-generated imagery, respectively. The uniformity of distribution was assessed using ***χ***^2^ tests: cue-induced imagery: ***χ***^2^(23) = 86.72, *p* < 0.001; self-generated imagery: ***χ***^2^(23) =1 31.51, *p* < 0.001. B. Same as a, but with results from Experiment 2: cue-induced imagery: ***χ***^2^(19) = 5.32, *p* = 0.503; self-generated imagery: ***χ***^2^(19) = 26.26, *p* < 0.001. C. Mean angular differences between initial probe orientations and final responses, in cue-induced (red) and self-generated (blue) imagery in Experiment 1. Colored circles indicate group mean (error bars denote ±1 SEM), gray lines indicated results from individual participants. Differences were averaged across trials for each participant and evaluated using one-sample t-test against 0: cue-induced imagery: *t*(23) = 0.75, *p* = 0.463; self-generated imagery: *t*(23) = 0.66, *p* = 0.518; D. Same as C, but with results from Experiment 2: cue-induced imagery: *t*(19) = 1.07, *p* = 0.297; self-generated imagery: *t*(19) =1.04, *p* = 0.31.

**Figure S3.**
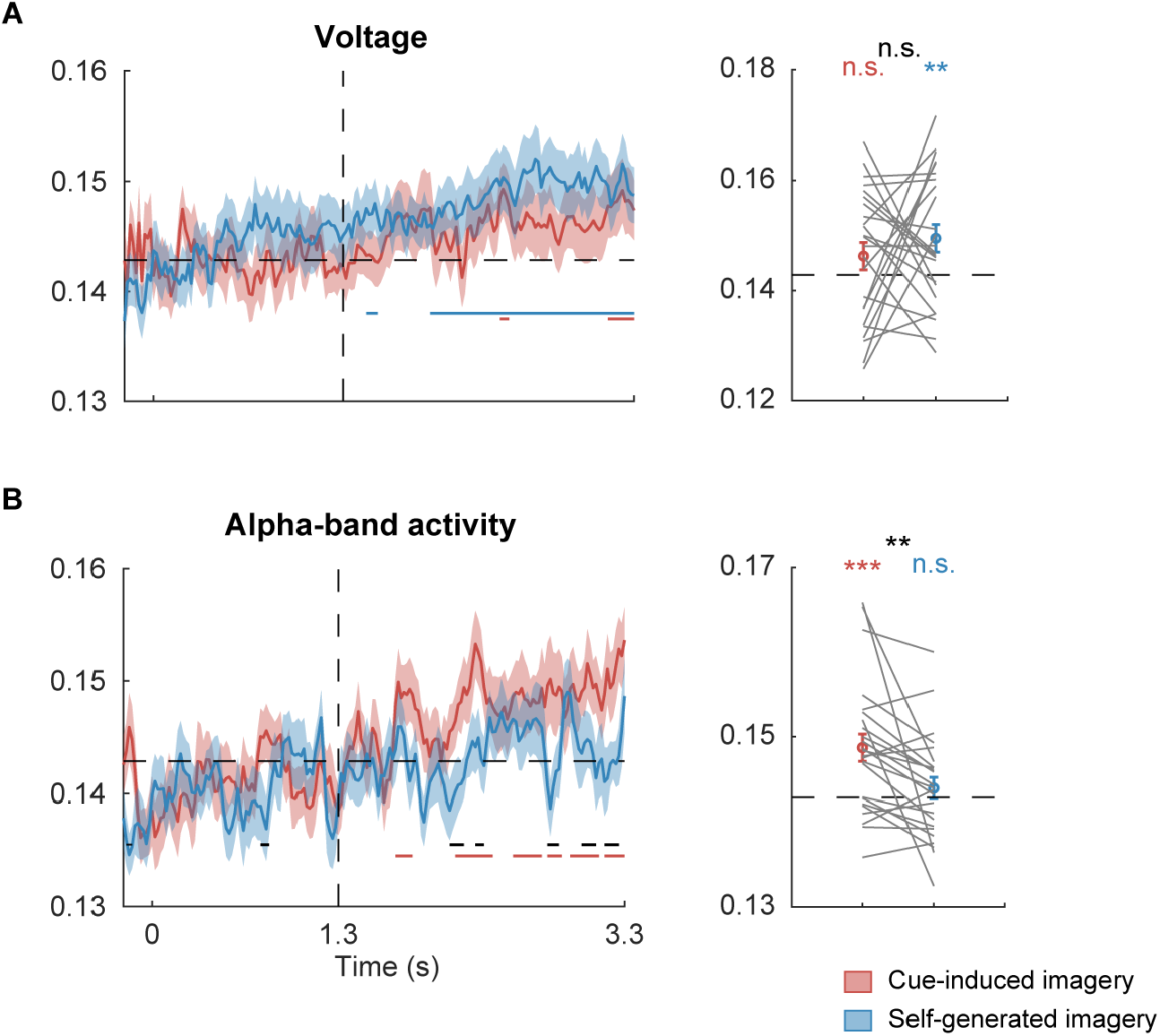
SVM decoding results in voltage and alpha-band activity from all electrodes. A. The left panel shows time course of decoding accuracy in cue-induced (red) and self-generated imagery (blue), from – 0.2 s prior to stimulus onset until end of delay. Y axis denotes decoding accuracy. Colored lines at the bottom denote significant time points of the corresponding condition, corrected for multiple comparisons using a cluster-based permutation method (*p* < 0.01). The vertical dashed line denotes onset of delay (at 1.3 s). The horizontal dashed line denotes chance level of 0.143 (i.e., 1/7). Shaded areas denote error bars (±1 SEM). The right panel shows decoding accuracies averaged over selected time periods of significance (2-3.3 s). Colored circles indicate group mean, error bars denote ±1 SEM, and gray lines indicated results from individual participants in cue-induced and self-generated imagery. Y axis represents decoding accuracy. Colored asterisks denote significance of the corresponding condition, and black asterisk denotes significance of difference between conditions. n.s., not significant, *: *p* < 0.05, **: *p* < 0.01, ***: *p* < 0.001. B. same as A, but with results from alpha-band data in all electrodes.

**Figure S4.**
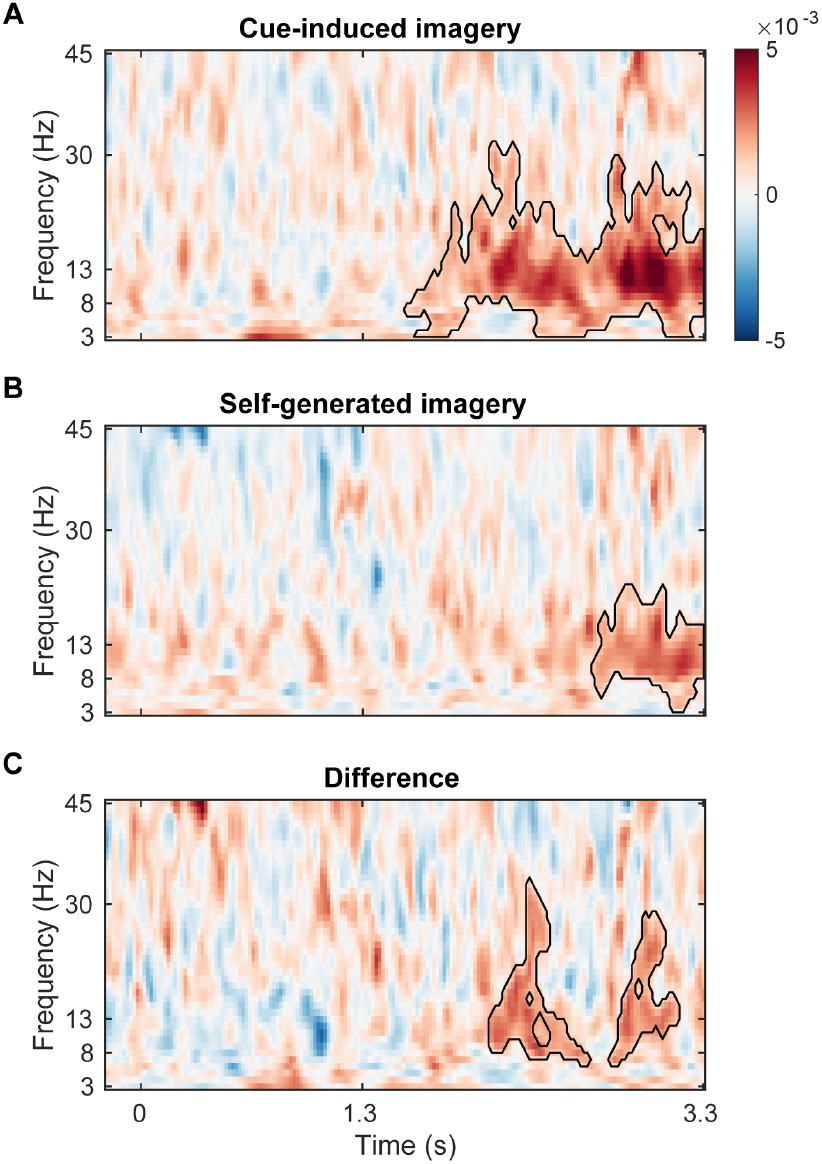
IEM results in frequencies from 3 to 45 Hz in EEG.Orientation representational strength, as reconstructed from power data in frequencies from 3 to 45 Hz in posterior electrodes of EEG, in cue-induced imagery (top), self-generated imagery (middle), and in difference between the two (bottom). X axis denotes time, and y axis denotes frequencies. Circled areas denote significant clusters determined using a cluster-based permutation method.

**Figure S5.**
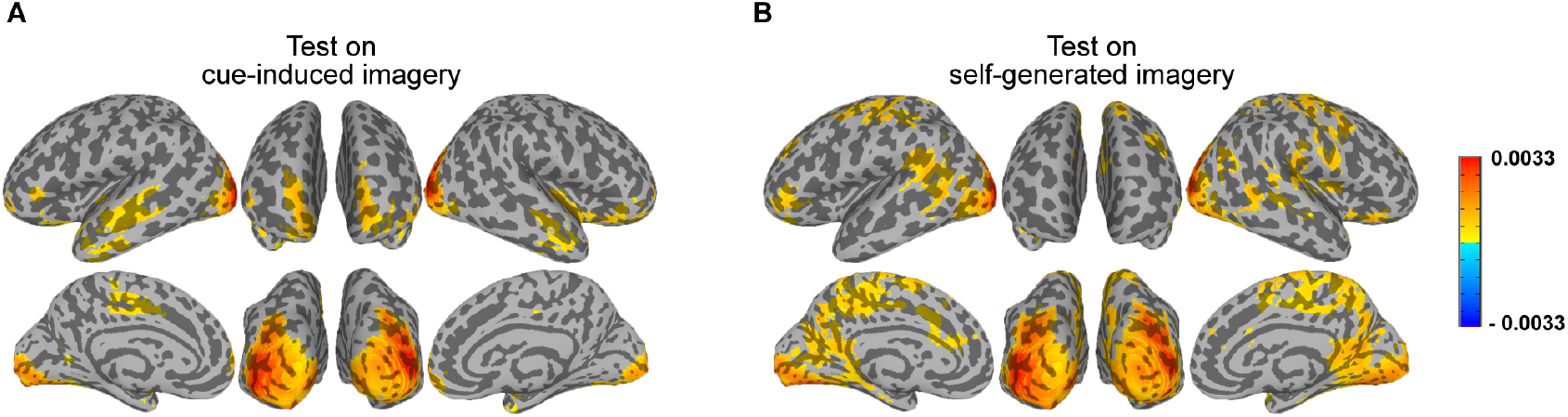
Whole-brain neural representations of imagery contents in late delay with a perception IEM. A. Searchlight parametric map of the strength of orientation representations in late delay (9-12 s; 6 s after retrocue) in cue-induced imagery, using an IEM trained from perception data. Colors on the cortical surface denote brain regions with significant orientation representations, corrected using a cluster-based permutation method (*p* < 0.01). For demonstration purposes, clusters were thresholded at 50 voxels. B. Same as A, but with results from self-generated imagery.

**Figure S6.**
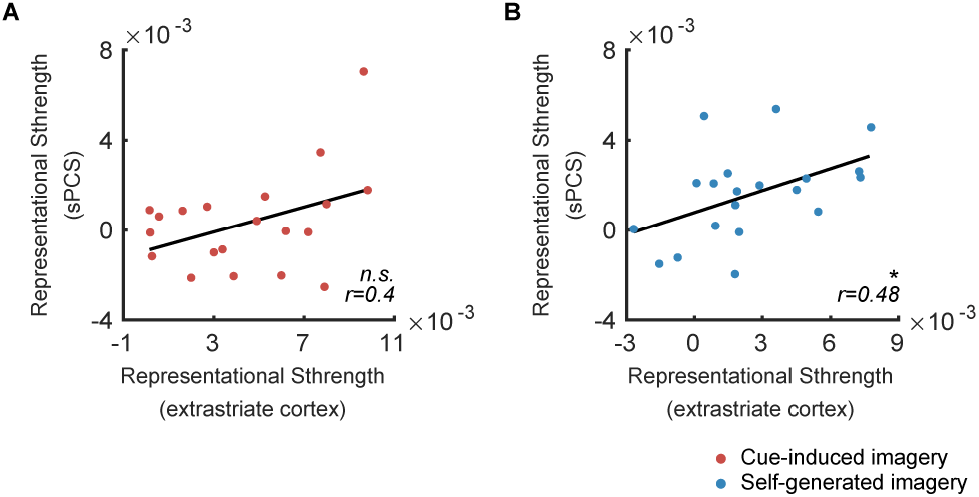
Correlations between the representational strength in right extrastriate cortex and sPCS. A. Pearson correlation between the representational strength in right extrastriate cortex and that in right sPCS in cue-induced imagery. B. Pearson correlation between the representational strength in right extrastriate cortex and that in right sPCS in self-generated imagery. Each dot represented individual participant. Representational strength in right extrastriate cortex (x axis) and representational strength in right sPCS (y axis) were averaged across trials for each participant. Black lines represented the best linear fit.

